# White eye-rings coevolved with diurnal behaviors as a trait enhancing visual appeal in rodents

**DOI:** 10.64898/2025.12.07.692862

**Authors:** Nguyen Hoang Khoi Le, Shou-Hsien Li, Chi-Cheng Chiu, Meng-Pin Weng, Shih-Kuo Chen, Ben-Yang Liao

## Abstract

Conspicuous colorations are widespread in animals, yet their adaptive functions remain unclear, particularly in mammals. White eye-rings (WER), the bright-colored pelage encircling eyes, are prevalence in rodents (Mammalia: Rodentia). Phylogenetic analyses across 601 rodents show that WER has repeatedly emerged with transitions to diurnalism and disappeared with reversions to nocturnalism during rodent evolution. To explore WER’s function, we performed behavioral tests on the Nile rat (*Arvicanthis niloticus*, diurnal with WER) and the house mouse (*Mus musculus*, nocturnal without WER), and found that WER enhances visual appeal (the ability to attract attention or preference), likely by accentuating facial bilateral symmetry. This effect is neither associated with sexual selection nor species recognition and was absent in mice. Thus, WER appears selectively favored in diurnal rodents capable of perceiving and responding to it, but is constrained in nocturnal species. These findings illustrate how ecological factors and shared sensory preferences shape conspicuous color patterns.

## Introduction

The diversity of animal coloration has attracted biologists to find its adaptive explanations for over a century and often leads to the advancement of evolutionary theories^1–3^. For instance, some conspicuous color patterns, such as dots and stripes, are recognized as sexual signals and have contributed to the development of sexual selection hypothesis^4–6^, though this has been subject to debate, especially in mammals where mate choice is frequently observed without apparent sexual dichromatism^7^. In recent years, significant advances have been made to understand molecular mechanisms and developmental pathways controlling the type, density, and distribution of skin and hair pigmentation in mammals^8–10^. Yet, the evolutionary drivers underlying the presence of conspicuous color patterns have not been fully understood.

Many terrestrial vertebrates have contrasting facial color patterns, which can be conspicuous and are often thought to serve functions such as camouflage, social or aposematic signaling, and regulation of physiological processes^2,3,11^. Notably, black patches around the eyes have been proposed to aid in concealment by disrupting facial outlines and obscuring the eyes^12^ or to enhance vision by reducing glare reflected into them^13,14^. In contrast, the adaptative meaning of white eye-rings (WER), the light-colored pelage encircling the eyes (Figure 1), remains poorly understood, despite the widespread occurrence of WER in many mammalian lineages.

**Figure 1.**
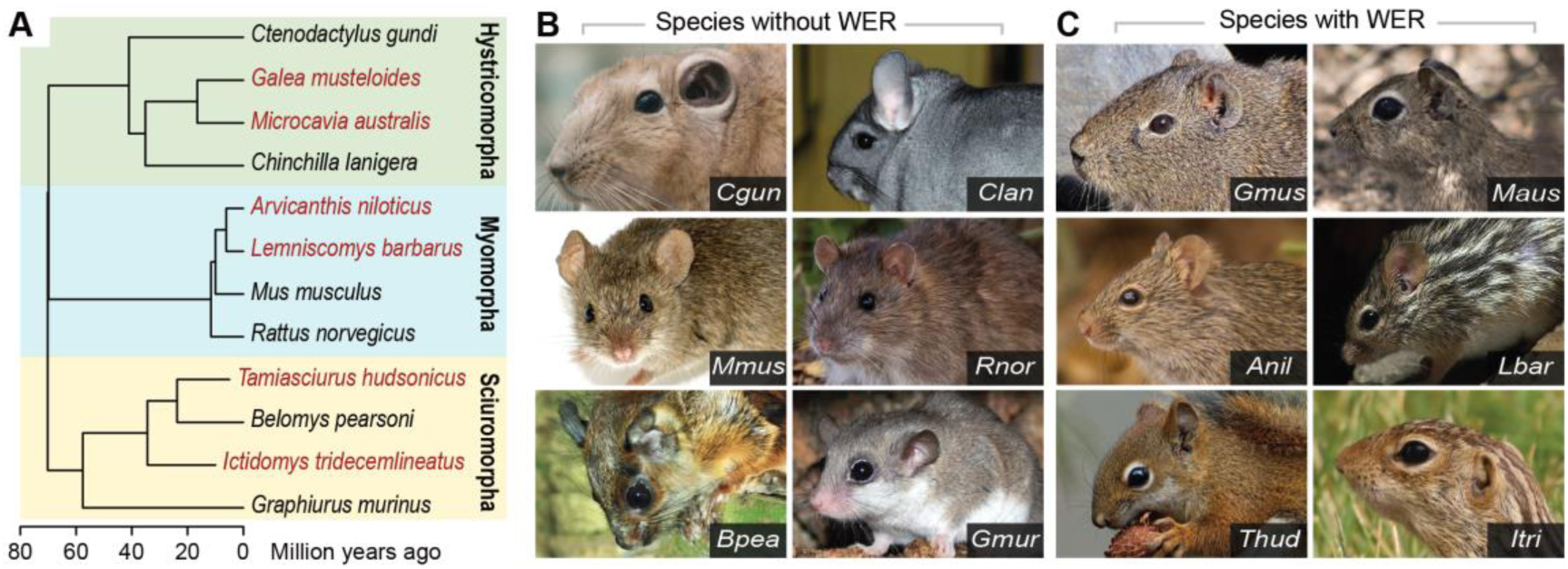
Examples of species with and without white eye-rings (WER) across three major lineages of Rodentia. (A) The phylogenetic tree with divergence times of 12 representative rodent species across three major suborders including Hystricomorpha, Myomorpha, and Sciuromorpha. Species with WER are highlighted in red color. (B) Examples of species without WER including *Ctenodactylus gundi* (*Cgun*), *Chinchilla lanigera* (*Clan*), *Mus musculus* (*Mmus*), *Rattus norvegicus* (*Rnor*), *Belomys pearsoni* (*Bpea*), and *Graphiurus murinus* (*Gmur*). (C) Examples of species with WER including *Galea musteloides* (*Gmus*), *Microcavia australis* (*Maus*), *Arvicanthis niloticus* (*Anil*), *Lemniscomys barbarus* (*Lbar*), *Tamiasciurus hudsonicus* (*Thud*), and *Ictidomys tridecemlineatus* (*Itri*). *Anil* and *Lbar* belong to the Arvicanthini tribe. All photos are licensed under CC-BY-SA 4.0 and CC-BY-NA 4.0.

Unlike black masks that conceal the eyes and improve vision, WER may instead accentuate these vulnerable regions and impair visual performance by increasing glare reflected directly into the eyes. Thus, unless the adaptive benefits of WER outweigh these disadvantages, they would not be expected to be so prevalent among species. Previous research suggested that WER might serve as distractive markings to help squirrels avoid predators^15^, or highlight important organs (i.e., preorbital glands) for communication among conspecifics in artiodactyls^16^. However, these proposed functions remain largely speculative and lack sufficient empirical support. Comparative studies in artiodactyls^17^ and canids^18^ have failed to establish any association between WER and social behaviors, casting doubt on its role in social communication and emphasizing the need for further investigation into the significance of this trait from a behavioral perspective.

Rodents are the most diverse and abundant mammalian order, comprising 2,693 extant species, which account for approximately 40.6% of all mammals^19,20^. Nevertheless, most attempts to explain the adaptation of rodent coloration have largely focused on non-facial regions^21–24^ despite the prevalence of WER in many rodent lineages (Figure 1). With considerable species richness, adaptability to diverse ecological niches, and frequent instances of phenotypic convergence or parallelism, rodents are well suited for studying evolution and adaptative significance of facial markings such as WER. To do so, we first examine the relationship between the presence of WER and various morphological, ecological, and social variables across rodent phylogeny. Our initial phylogenetic comparative analysis reveals that the presence of WER is tightly linked to diurnal activity. Based on this result and the fact that diurnal vertebrates have increased visual acuity than nocturnal^25^, we then hypothesized that rodents transitioning from a nocturnal to a diurnal temporal niche develop the capacity to perceive WER as social and/or sexual signals. To test this, we performed a series of behavioral experiments on the Nile rat (*Arvicanthis niloticus*) and the house mouse (*Mus musculus*) using three-chambered behavioral tests. After diverging from the lineage of the house mouse (*Mus musculus*), the Nile rat switched its temporal niche from nocturalism to strict diurnalism^26^, evolved more acute vision^27,28^, and acquired conspicuous WER (Figure 1C). This study seeks to uncover the evolutionary drivers of the emergence of light-colored facial markings in animals.

## Results and Discussion

### Distribution of white eye-rings and circadian activity patterns across rodent phylogeny

In this study, we compiled phenotypic variables (Supplementary Table 1) of 601 species (representing 22.3% of Rodentia) across 5 suborders: Anomaluromorpha (n = 3), Castorimorpha (n = 53), Hystricomorpha (n = 80), Myomorpha (n = 289), and Sciuromorpha (n = 176) (phylogeny: Supplementary Data 1; phenotypes: Supplementary Data 2). Among these, we found WER existed in 145 species from Hystricomorpha, Myomorpha, and Sciuromorpha had WER, but absent in species from Castorimorpha and Anomaluromorpha (Supplementary Table 2). Specifically, 116 out of 168 sampled species from the family Sciuridae of Sciuromorpha had WER, whereas only 20 out of 289 sampled species within Myomorpha (mainly in the Muridae family) and 9 out of 80 sampled species within Hystricomorpha (mainly in the Caviidae family) exhibited this trait.

Squirrels (Sciuridae species) are known to be predominantly diurnal, while murids (Muridae species, including rats, mice, voles) are predominantly nocturnal^29^. Our compiled dataset supported existing trends, showing that diurnal species were prevalent in squirrels (151 out of 176 sampled species) and comparatively rare in murids (29 out 289 sampled species) (Supplementary Table 2). Among murids, the Arvicanthini tribe (suborder *Myomorpha*, family *Muridae*) is an exception, with many species exhibiting a diurnal lifestyle^29^. Interestingly, many of these Arvicanthini species also display WER (Figure 1). These documented trends, along with our finding on the prevalence of WER, suggest a potential association between WER and diurnalism.

To investigate this anticipated association, we inferred the evolutionary history and ancestral states of WER or circadian activity patterns across rodent phylogeny through stochastic character mapping^30^ (see Materials and Methods). It was revealed that the most recent common ancestor (MRCA) of rodents was likely nocturnal and lacked WER (Figure 2 and Supplementary Table 3). This result aligns with an earlier study examining over 2400 species from all extant mammalian orders^31^. More importantly, we found that the emergence of WER independently occurred across the three major rodent suborders, coinciding with the evolutionary transitions to a diurnal activity from ancestrally nocturnal rodents. The earliest appearance of WER occurred approximately 34.6 million years ago (MYA) since the divergence of the squirrel lineage (suborder Sciuromorpha, family Sciuridae) (Supplementary Table 3). Similar events were found in the MRCA of a branch in the Arvicanthini tribe around 7.35 MYA and in the MRCA of the family Caviidae around 11.9 MYA, where species in these lineages acquired both WER and diurnal activity. Interestingly, an event occurred in the Pteromyini tribe (suborder Sciuromorpha, family Sciuridae) around 16.8 MYA where WER disappeared and nocturnalism evolved.

**Figure 2.**
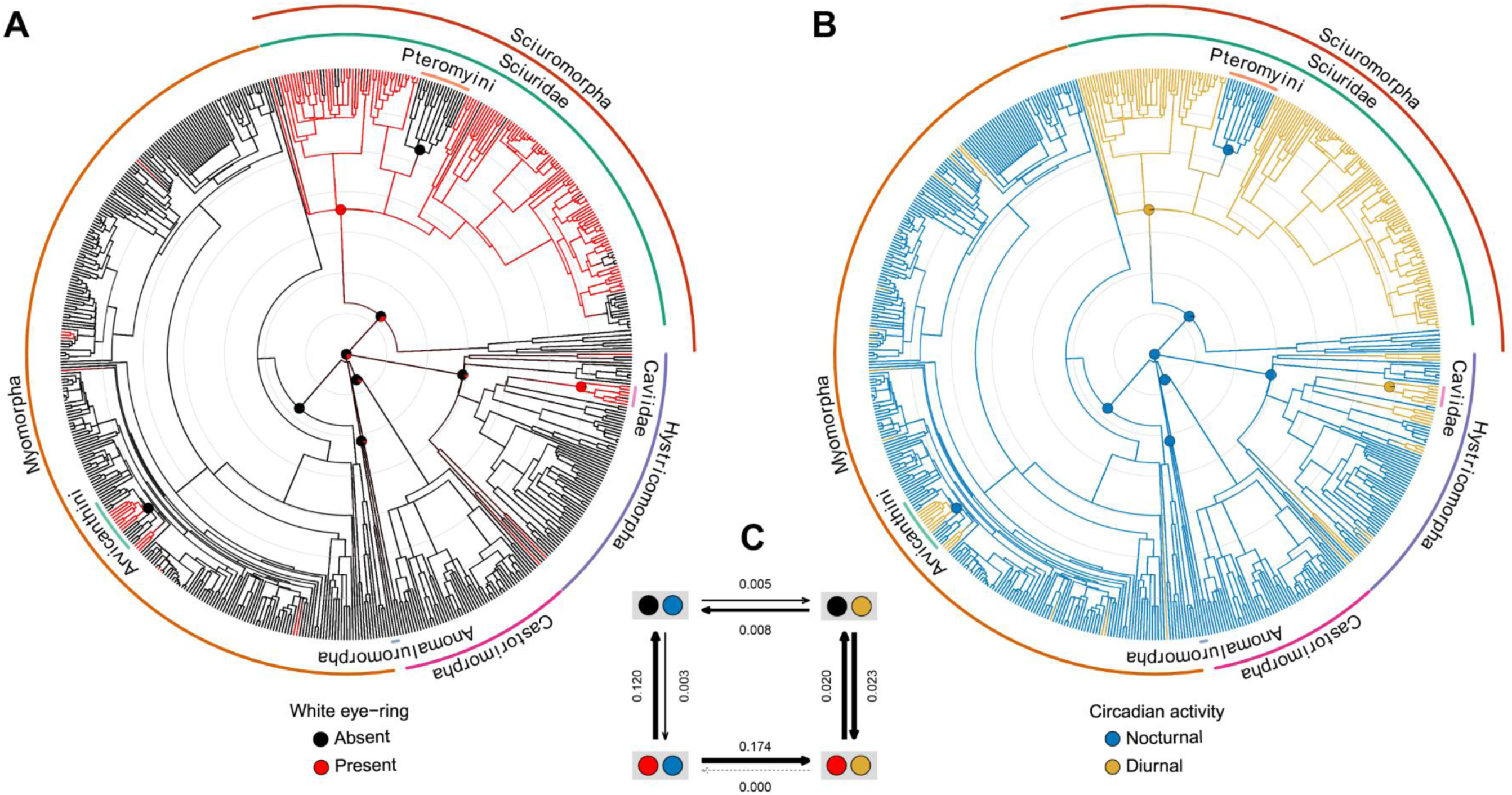
The correlated evolution of white eye-ring (WER) and circadian activity patterns across rodent phylogeny. (A-B) The phylogenetic tree with divergence time and reconstructed ancestral states of WER (A) and circadian activity (B). Each gray ring represents a duration of 10 million years since the divergence of Rodentia order ∼70 MYA (the center node). The pie charts indicate the posterior probability (PP) of the ancestral states at key nodes (for a complete result, see Supplementary Figure 1; for PP of the highlighted nodes, see Supplementary Table 3). (C) Evolutionary transition rates of WER and circadian activity patterns in rodent species. The transition rates were calculated by assuming that the evolution of these two traits is interdependent. Thicker arrows denote higher transition rates, whereas dashed arrows represent the absence of observed transitions.

### Correlated evolution of white eye-rings and circadian activity

Closely related species tend to exhibit similar traits due to shared ancestry (i.e. the presence of phylogenetic signal), which can lead to false positive determinations of correlated evolution among traits^32^. To verify the evolutionary relatedness of WER and circadian activity patterns, we tested for the strength of phylogenetic signal in both traits using *D*-statistics^33^ (see Materials and Methods). *D*-statistics assesses whether the trait of interest evolves under phylogenetic randomness model (*D* ≈ 1, no phylogenetic signal) or Brownian motion model (*D* ≈ 0, strong phylogenetic signal) along the phylogeny. The estimated *D* values of both WER and circadian activity patterns were not significantly different from 0 (WER: *D* = -0.243, *P* = 0.887; circadian activity pattern: *D* = -0.316, *P* = 0.953), suggesting strong phylogenetic signals in the analyses of both traits (Supplementary Table 4). Therefore, approaches that account for phylogenetic signals among taxa should be used to test whether the evolution of WER and diurnalism are correlated.

Our first approach was the Pagel’s 1994 method, which models how the rate of change in one character can influence the rate of change in the other^34^. The result showed that the dependent model, where the transition rates of WER and circadian activity patterns are interdependent during evolution, had a significantly better fit than the independent model (ΔAIC = 58.335, *P* < 10^-12^; AIC, Akaike information criterion) (Supplementary Table 5). To further investigate the directionality of the evolutionary relationship between WER and circadian activity, we compared different dependent models: (1) “dependent WER model”, where WER transitions depend on changes in circadian activity; (2) “dependent circadian activity model”, where circadian transitions depend on WER; and (3) “interdependent model”, where transitions in both traits are mutually dependent. The interdependent model was found to have the strongest support, indicated by its highest Akaike weight (AICw) (0.972) (Supplementary Table 5). Under this interdependent model, the estimated transition rates suggest that, during rodent evolution, diurnalism promoted the emergence of WER (00.23 - 0.020 = 0.003), whereas nocturnalism favored the loss of WER (0.120 - 0.003 = 0.117) (Figure 2C). Likewise, the presence of WER promoted shifts from nocturnalism to diurnalism (0.174 - 0 = 0.174), while its absence facilitated transitions of circadian activity in the opposite direction (0.008 – 0.005 = 0.003) (Figure 2C). The estimated rates of transitions from the “nocturnalism, with WER” may be prone to error due to the rarity of species in this state (only 4 out of 601); therefore, our emphasis is placed on the direction rather than the absolute magnitude of these transition rate estimates.

The phylogenetic logistic regression for binary traits^35^ was employed as the second approach to identify the regression (rather than correlation as tested by Pagel’s method) between two traits. The result showed that the presence of WER is more strongly dependent on diurnalism (log odds ratio = 4.534, standard error (SE) = 0.962, *P* < 10^-5^) than diurnalism is on WER (log odds ratio = 1.991, SE = 0.463, *P* < 10^-4^), with both associations statistically significant (Supplementary Table 6). This result is not inconsistent with the results from Pagel’s test, as regression-based method tests the statistical association between trait values at the tips instead of historical contingency in evolutionary transitions like in Pagel’s method. Although the detailed causal relationship between WER and diurnalism remains to be revealed due to a discrepancy in selecting the best fitting model between the two methods employed and mentioned above, the consistent identification of diurnalism as a potential driver for the emergence of WER, after correcting for phylogeny, warrants further hypothesis-driven investigation.

Diurnal rodents have evolved with more advanced visual systems compared to their nocturnal relatives^25,27,36,37^. Previous comparative studies further suggested that the shift to a diurnal niche, accompanied by enlarged visual regions of the brain and enhanced visual acuity, could lead to the degeneration of the olfactory system as an adaptation to saving energy cost ^38,39^. Although the precise impact of this vision-olfaction trade-off on social behaviors in rodents remains unclear, we hypothesize that diurnal rodents with acute vision may utilize conspicuous pelage markings (e.g. WER) for visual signaling which facilitates social communication, thereby potentially compensating for the social interactions once mediated by olfactory cues in their nocturnal ancestors. Conversely, the absence of WER in nocturnal rodents may reflect constraints imposed by low-light environments, where limited visual capabilities diminish the selective advantage of such traits. However, it is critical to further confirm that the association between WER and diurnalism is not confounded by other ecological variables, and that WER directly influences behaviors.

### No association between white eye-rings and other body marks, habitat preference, or social organization

The association between WER and diurnalism may result from the confounding effects of other variables. For example, WER could be derived from white markings on other body parts such as white stripes or spots extending from the back and should not be considered as an independent trait. Moreover, squirrels living in forests are known to have white body marks primarily to disrupt body outline for concealment or distraction^15,40^. These forest-dwelling species also tend to be diurnal^29^, which may explain the observed link between WER and diurnal behavior. Additionally, the yellow-bellied marmot *Marmota flaviventris*, a diurnal rodent that has evolved a moderately complex, kin-based social organization that includes harem-polygynous mating^41^, displays individually variable white facial markings^42^, suggesting that social structure may also mediate the relationship between WER and diurnalism. Thus, to determine whether potential confounding factors affect the WER-diurnalism association, we further analyzed pelage color data for non-WER white markings, along with socioecological variables, including habitat preference and social organization.

Our analysis revealed strong phylogenetic signals in both non-WER markings (*D* = 0.139) and habitat preference (forest: *D* = 0.165; desert: *D* = 0.319, but not grassland) (Supplementary Table 4), which emphasized the importance of controlling for phylogeny when examining the correlation between WER versus any of these two traits. We therefore again conducted phylogenetic logistic regression, and the results showed no association between WER and non-WER markings (log odds ratio = 0.302, SE = 0.263, *P* = 0.249) (Supplementary Table 7). Similarly, although body stripe patterns in rodents are thought to help them blend in with grassland or tree trunks for concealment^21^, the phylogenetic logistic regression showed no association between the presence of WER and any habitat type, including forest (log odds ratio = 0.053, SE = 0.153, *P* = 0.728), grassland (log odds ratio = 0.201, SE = 0.154, *P* = 0.192), and desert (log odds ratio = -0.160, SE = 0.225, *P* = 0.477) (Supplementary Table 7). This lack of association may be explained by the independent evolution of WER and non-WER white markings, or because, across rodents, concealment through substrate color matching in different micro-habitats is primarily achieved through whole-body coloration or body patterns^43,44^, rather than by facial markings such as WER.

Although social organization did not exhibit strong phylogenetic signal (*D* = 0.495, Supplementary Table 4), we applied phylogenetic logistic regression for consistency with other analyses. No correlated evolution between WER and social organization was identified (Supplementary Table 7; log odds ratio = -0.365, SE = 0.208, *P* = 0.079), despite the presence of diurnal social rodents with white facial markings (e.g., *M. flaviventris*). The above-mentioned results indicated that the correlated evolution of WER and diurnalism is unlikely to have resulted from potential confounding factors that were tested.

### Preference for individuals with WER over those without in Nile rats

Why did diurnal rodents evolve WER? To explore this question, we studied the behavior of the Nile rat (*Arvicanthis niloticus*), a member of the Arvicanthini tribe, which consists of several closely related African murids with conspicuous WER and recently acquired diurnal activity (Figure 2 and Supplementary Figure 2). We hypothesized that WER serves as a visual signal to promote intraspecies communication from a distance in diurnal rodents. To test this hypothesis, we evaluated WER’s influence on the behavior of the Nile rats using the three-chambered test (Figure 3A), which is a common method to assess animal social behaviors such as sociability and signal preference^45^.

**Figure 3.**
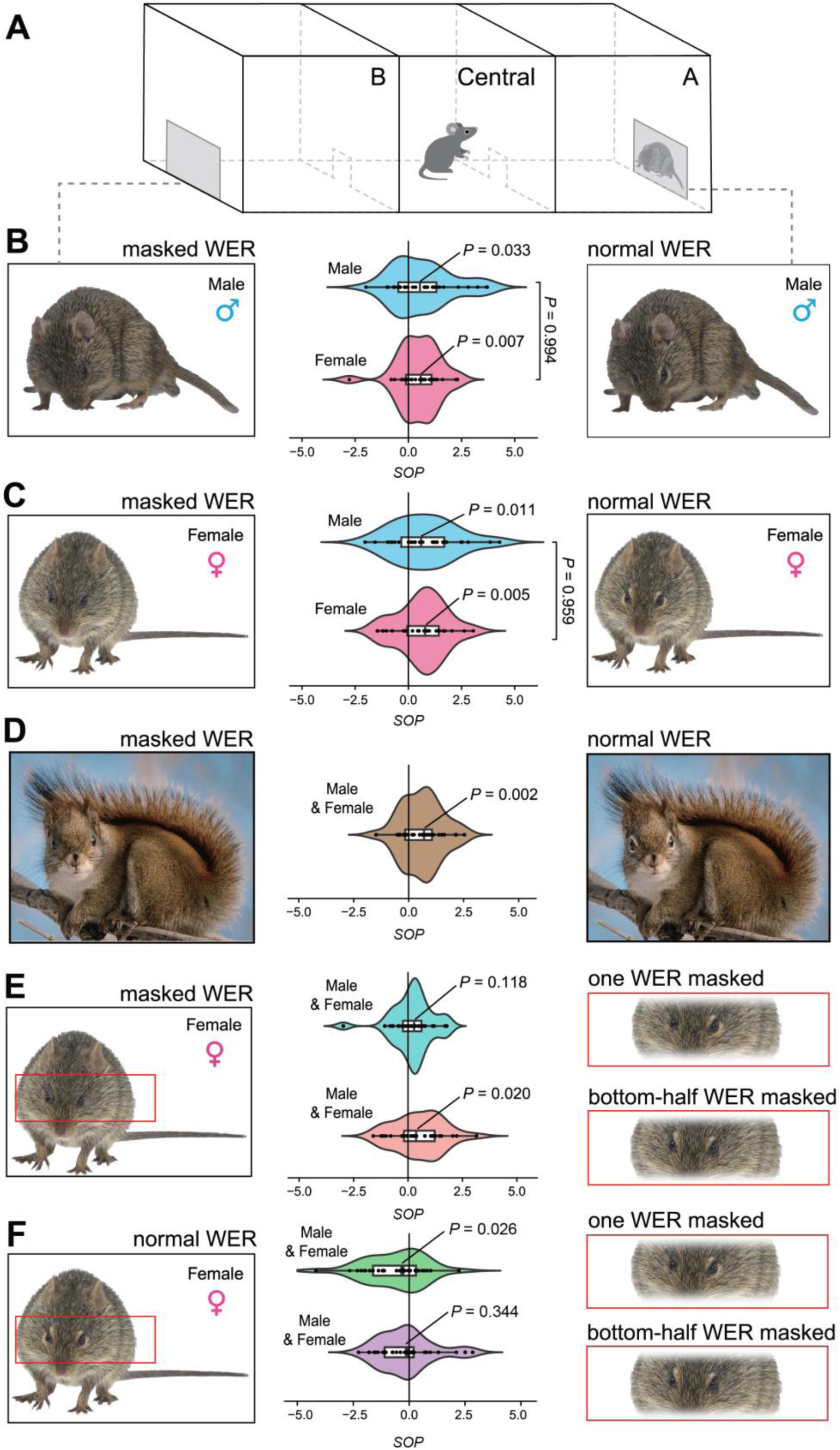
Preference for white eye-rings (WER) in the Nile rats. (A) Schematic of the three-chamber sociability test showing chamber A (right), the central chamber (middle), and chamber B (left). (B–C) Violin plots of strength of preference (*SOP*) in male (n = 30; light blue) and female (n = 30; pink) Nile rats for images of a male (B) and a female (C) Nile rat displaying WER, compared to the same images with masked WER. (D–F) Violin plots of *SOP* in male and female Nile rats (n = 30; 15 males and 15 females) toward: (D) an image of an American red squirrel (*T. hudsonicus*) displaying WER versus the same image with masked WER (brown); (E) a Nile rat image with an asymmetrical WER (cyan) or a symmetrical pattern (coral) versus the masked WER image; and (F) Novel stimuli (asymmetrical WER: green; symmetrical pattern: purple) versus non-novel stimuli (normal WER). For all violin plots, the internal box indicates the lower quartile, median, and upper quartile; whiskers represent quartile ranges. *P*-values for median *SOP* were obtained using a one-sample Wilcoxon signed-rank test (null hypothesis: *SOP* = 0). Between-group comparisons were performed using two-sided Mann–Whitney *U* tests (null hypothesis: equal medians). Photos of *T. hudsonicus* are licensed under CC-BY-NC 4.0.

Given the limited evidence on the capacity of diurnal rodents to discern conspecific sex from two-dimensional images, we employed two treatments, each displaying two images (original vs. photo-edited) of an individual from one sex (male image: Figure 3B; female image: Figure 3C). Both male and female animals were included in each treatment to evaluate the potential visual preference for WER. A preference, or lack thereof, for WER in the test would support or reject the influence of WER in social communication. Testing animals of both sexes could provide further insights into the type of social communication WER is involved in. If a preference for WER is verified and this preference is linked to sexual selection, one sex is expected to show a stronger preference than the other. However, if both sexes are equally attracted to WER, it suggests that WER primarily serves as a visual cue for communication in non-mating contexts. In the first treatment, when presented with images of a male Nile rat, both male and female subjects showed a significantly stronger preference in exploring the chamber containing the image of a rat with its WER (Chamber A, as shown in Figure 3A) over the chamber containing the image of the same rat with its WER masked (Chamber B, as shown in Figure 3A), as indicated by their median *SOP* values (see Materials and Methods) being greater than 0 (in male: *P* = 0.033, n = 30; in female: *P* = 0.007, n = 30; Figure 3B). Moreover, there was no significant difference in *SOP* between male and female animals (*P* = 0.994; Figure 3B). In the second treatment, when exposed to images of a female Nile rat, again both male and female subjects showed significantly stronger preference for chamber A over chamber B (in male: *P* = 0.011, n = 30; in female: *P* = 0.005, n = 30; Figure 3C), while *SOP* did not differ significantly between males and females (*P* = 0.959; Figure 3C). This finding suggests that WER influences social interactions, but its role is unrelated to sexual selection in Nile rats, consistent with the absence of sexual dichromatism in their pelage color patterns.

### Preference for symmetrical facial marks over nonsymmetrical ones in Nile rats

Why and how does WER enhance visual appeal in diurnal rodents? Animals may have commonly shared visual preferences for color patterns that resemble specific stimuli, such as diet or biological symmetry, which drive their phenotypic evolution^46,47^. To investigate whether such a visual preference exists in the Nile rats, we expanded our three-chambered experiments by replacing images of the Nile rats with those of an American red squirrel, *Tamiasciurus hudsonicus*, which also has striking WER, though in a form that differs from the Nile rat’s WER (i.e. brighter color of WER, different facial and body coloration, and different animal shape). This test aimed to determine whether WER enhances species recognition. Given that no behavioral differences were observed between male and female Nile rats (Figure 3B-C), animals of both sexes were pooled for the test. Interestingly, our result revealed that the tested Nile rats still showed significantly higher interest in chamber A (containing the image of a normal squirrel) than in chamber B (containing the image of the same individual but with masked WER) (*P* = 0.002, n = 30; Figure 3D). Since the Nile rats are native in Africa, it is highly unlikely for them to encounter American red squirrels in the wild. The Nile rats’ visual preference for the American red squirrel’s WER indicates that their preference for WER is not tied to recognizing their species and may have existed before their ancestors acquired WER.

In Bilateria, bilateral symmetry is widely considered as a visual cue to attractiveness^47–49^, and body color patterns potentially amplifies this effect^50,51^. To determine whether the Nile rat’s visual preference for WER stems from its bilateral symmetry or donut-like shape, we analyzed the animal’s differential responses to modified images: one with a rat’s WER masked on one side (asymmetry, top of Figure 3E) and another with the bottom half of both sides of WER masked (symmetry, bottom of Figure 3E) (see Materials and Methods). The results showed that the animals did not display a statistically significant preference for the intact, full-sized asymmetrical WER over the masked WER (*P* = 0.118, n = 30; Figure 3E, top). However, they were significantly more drawn to the half but symmetrical WER (*P* = 0.020, n = 30; Figure 3E, bottom). Given that murids tend to explore novel stimuli less than familiar ones^52,53^, the symmetrical half-WER pattern likely represented a novel visual configuration. To test whether novelty alone accounted for this preference, we compared responses to two symmetrical conditions: fully masked (familiar) and bottom-half masked (novel) (Figure 3F, bottom). No preference was observed, suggesting novelty does not drive the effect (*P* = 0.344, n = 30, Figure 3F). Together with the results from the squirrel image test (both novel stimuli; Figure 3D), these findings support the interpretation that visual preference for WER in Nile rats is driven by bilateral symmetry rather than novelty or shape of the facial mark.

Previously, facial color patches have been proposed as important factors in intraspecies communication, such as sexual selection, conspecific recognition, and social interaction in mammals, though empirical evidence is scarce^7,12,54^. Our behavioral experiment revealed that Nile rats are attracted to symmetrical color pattern on the face, which WER accentuates, regardless of the sex of receivers and species of the stimuli (the still images). This visual preference for bilateral symmetry is likely innate and shared across species, as it appears unrelated to species recognition. Therefore, the WER observed in other diurnal rodents may have arisen for the same reason.

### Lack of visual preference for WER in house mice

To examine whether preference for WER also occurs in nocturnal rodents, we repeated the three-chambered behavioral experiment on the C57BL/6 (B6) strain of the house mouse (*Mus musculus*). The B6 mouse strain provides an appropriate comparison to the Nile rat for several reasons: it has a uniformly black coat lacking conspicuous markings, exhibits relatively good vision among inbred mouse lines^55^, and has been reported to use visual cues in social interactions^56^. Test images consisted of wild house mice with a natural agouti coat, either unaltered or digitally modified to display WER (Figure 4). Using these images controlled for the novelty of visual stimuli, as the tested animals had never encountered individuals with or without WER on an agouti coat background prior to the experiment. In contrast to Nile rats, neither male nor female B6 mice showed a significant preference for the image with WER (in male: *P* = 0.349, n = 20; in female: *P* = 0.154, n = 20; Figure 4). These results indicate that visual attraction to WER is absent in nocturnal rodents. Whether this difference reflects the loss of an ancestral visual bias in nocturnal lineages, the gain of such a preference in diurnal rodents, or a combination of both evolutionary processes remains to be determined.

**Figure 4.**
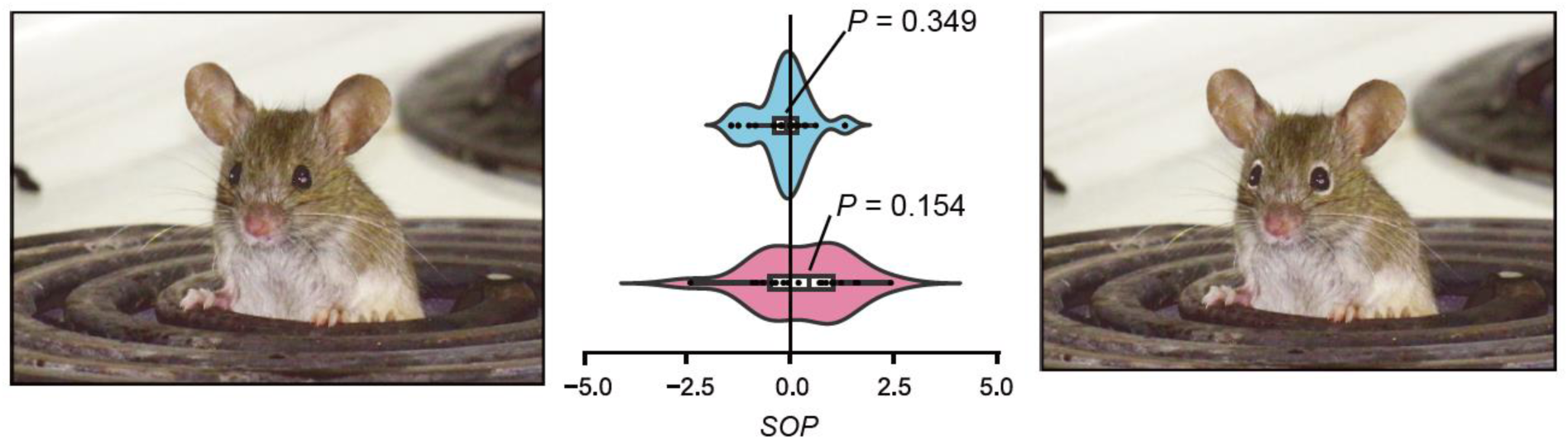
No preference for white eye-rings (WER) in the B6 house mice. The experimental setup is the same as in Figure 3A. Violin plots show *SOP* in male (n = 20; light blue) and female (n = 20; pink) B6 mice for images of a wild house mouse that are unaltered (left) versus the same images with digitally added WER (right). Internal boxes indicate the lower quartile, median, and upper quartile; whiskers show quartile ranges. *P*-values for median *SOP* were obtained using a one-sample Wilcoxon signed-rank test (null hypothesis: *SOP* = 0). The unmodified photo is licensed under CC-BY-NC 4.0.

### Concluding remarks

To investigate the adaptive role of WER from a macroevolutionary perspective, this study revealed a strong association between the evolution of WER and the evolutionary shifts to diurnal activity from the ancestrally nocturnal states over the course of rodent evolution. This pattern suggests that WER emerged predominantly in diurnal rodents, while remaining evolutionarily constrained in nocturnal species. The correlated evolution between WER and diurnalism is unlikely to have resulted from potential confounding factors, including other non-WER markings, habitat preference, and social organization. This repeatability and specificity indicate the importance of WER in diurnal rodents and highlight how environmental factors and sensory physiology can interact to influence the emergence, maintenance, and loss of novel pelage color patterns in mammals.

Our behavioral experiment further provided the empirical evidence suggesting that a preference for bilateral symmetry, unrelated with sexual selection and species recognition, has driven the evolution of facial color patterns in murine, a lineage traditionally considered reliant on olfaction rather than vision for social communication. In species with sufficient visual capability (e.g., the Nile rat), visual cues that enhance bilateral symmetry may serve as a form of sensory exploitation^57^, drawing attention and increasing the likelihood of conspecific social interactions. Although the direct fitness consequences of WER remain to be tested, it is known that house mice can detect stressed or diseased individuals through pheromones and odor cues, thereby minimizing social contact with infected individuals^58^. Given the high ectoparasite burden in the wild Nile rats and their documented vision-olfaction trade-off, conspicuous and symmetrical WER might function as a visual cue indicating parasite-free status, thereby facilitating safer social interactions prior to close-range contact. This signaling function is likely ineffective in dim-light environments. As a result, WER does not serve the same functions in nocturnal rodents as in diurnal species and may be constrained in nocturnal lineages, consistent with our observation that house mice show no preference for WER.

Interestingly, we found that the Nile rat’s preference for symmetry extends beyond conspecific stimuli. It remains unclear whether rodent predators show a similar type of preference for WER, potentially increasing predation risk for WER-bearing rodents. However, this risk is likely minimal, as predators rarely engage in face-to-face encounters with their prey, unless WER compromises the concealment provided by dark pelage in nocturnal rodents. It is also possible that WER is constrained in nocturnal rodents simply due to the absence of selective advantages in low-light environments. Future research should examine whether similar behavioral patterns exist in other rodent species, both diurnal and nocturnal, with and without WER, to validate the generality of our hypothesis. Additionally, it should investigate how potential nocturnal predators respond to WER to clarify the possible evolutionary costs linked to this trait. Exploring these ecological dimensions will deepen our understanding of WER’s role and cost in mammalian adaptation to temporal niche shifts.

## Materials and Methods

### Phenotypic data collection

Our dataset consists of phenotypic traits for 601 rodent species (Supplementary Data 2). The traits included in the dataset are detailed as follows: The coat-related traits for each species were determined based on images found in published literature or images deposited in public domains. For each rodent species, the presence or absence of WER was determined using ImageJ2 software^59^ for image thresholding to define color pattern boundaries with minimized human visual bias. In addition, each species was characterized if it possesses additional white markings (non-WER markings) such as facial masks or stripes, body spots, blotches or stripes, black-and-white quills, or tails.

The compiled data of socioecological variables consist of circadian activity patterns, habitat preference, and social organization (Supplementary Table 1; Supplementary Data 2). The data of the circadian activity patterns were obtained from the literature^29,60–64^, where animals are generally classified as diurnal (primarily active during daylight), nocturnal (primarily active at night), crepuscular (active during twilight), or cathemeral (active at any time of the day, with activity influenced by environmental factors). To simplify the statistical analyses, we excluded crepuscular and cathemeral species from our database, focusing on those classified as either diurnal or nocturnal species. The information on habitat preference was retrieved from the literature^62,63^ and the International Union for Conservation of Nature (IUCN) Red List database (https://www.iucnredlist.org/). We classified habitats based on vegetation levels into three categories: (1) forest, (2) grassland, and (3) desert. For social organization, species were categorized as either “solitary” or “group-living” based on published data^61,65^.

### Testing for phylogenetic signal

The study of trait evolution across multiple species requires considering the phylogeny of the species being examined. We first obtained the rooted phylogeny with estimated divergence time for the 601 Rodentia species from TimeTree 5 (https://timetree.org/)^66^. It should be noted that certain parts of the downloaded phylogeny were inconsistent with updated knowledge, and were manually corrected based on the literature ^67^ (Note in Supplementary Data 1). Additionally, the downloaded divergence times are median estimates from multiple studies and do not include information on their associated uncertainty. These uncertainties in both the phylogeny and the estimated divergence times could limit the reliability of our phylogeny-based analyses. To determine the strength of the phylogenetic signal of the focal traits, we used the phylogenetic *D*-statistics approach implemented in *phylo.d* function in R package ‘caper’ version 1.0.3. In general, the observed *D* value (the observed phylogenetic signal), which is the sum of changes between two estimated nodal values of a binary trait in the phylogeny, is compared with the alternative *D* values generated from 1000 simulations under each of the two models: Brownian motion (strong phylogenetic signal) and phylogenetic randomness (no phylogenetic signal)^33^. Both observed and simulated *D* values were then scaled to 0 (Brownian motion) and 1 (random distribution), using the means of two simulated datasets for calibration. A nonsignificant *D*-statistic from 0 suggests that species’ traits evolve following the Brownian motion model, while a value of 1 indicates random trait evolution independent of phylogeny.

### Ancestral state reconstruction

To explore the evolution of WER and circadian activity, we reconstructed the ancestral state of the two focal traits using ‘phytools’ package^68^. We first evaluated the fit of two evolutionary models, equal rate (ER) and all rates different (ARD), using *fitMk* function. Likelihood ratio tests suggested that the ARD model provided a significant better fit for both traits (*P* < 0.05). Based on this, we performed stochastic character mapping^30^, using the *make.simmap* function. A total of 1000 stochastic character maps were generated, with the transition matrix *Q* of the ARD model sampled from its posterior distribution every 10 generations (Q = “mcmc”, samplefreq = 10). The prior distribution on the root node of the tree was set as the empirical mean of the transition matrix (use.empirical = TRUE). Using the *summary* function, we calculated the posterior probabilities of the states at each internal node.

### Correlated evolution analyses

To examine the evolutionary dependency between the two focal traits (e.g., WER vs. circadian activity pattern), we used two approaches: Pagel’s correlation method proposed in 1994^34^ and phylogenetic logistic regression^35^. First, Pagel’s method, implemented in *fitPagel* function of R package ‘phytools’ version 2.0-3^68^, was used to fit the evolution of two binary characters under both independent and dependent models. The independent model assumes the two traits evolve independently from one another across phylogeny. In the dependent models, three assumptions were tested: (1) the transition rates of WER depend on the state of circadian activity, (2) the transition rates of circadian activity depend on the state of WER, and (3) the transition rates of both traits are interdependent. Likelihood ratios of independent and dependent models were compared using χ^2^ test. To infer the directionality of trait dependence, we identified the best-supported causal models based on Akaike weights.

For the second approach, we performed a phylogenetic logistic regression using *phyloglm* function in R package ‘phylolm’ version 2.6.2^69^. This analysis employed the optimization method “logistic_MPLE”, which maximizes the penalized likelihood of the logistic regression. We performed 2000 bootstrap replicates to compute the confident intervals and *P*-values of the estimates.

### Animals and three-chambered apparatus

All animal breeding and experimental procedures were approved by the Institutional Animal Care and Use Committee of the National Health Research Institutes (NHRI) (Approval No. IACUC-111103-AC1-M2-A). Male and female Nile rats were descendants of two breeding pairs from the colonies maintained by the KC Hayes Laboratory at Brandeis University. The B6 mice were obtained from the National Center for Biomodels, Taiwan. The animals were housed at the Laboratory Animal Center of National Health Research Institutes under 12h/12 h (B6 mice) and 11h/13h (Nile rats) light/dark cycles and were provided food (B6 mice: MFG, Oriental Yeast Co. Ltd., Tokyo, Japan; Nile rats: LabDiet 5326, LabDiet Ltd., St. Louis, MO, USA) and water at libitum.

A custom-designed apparatus to conduct three-chambered sociability test was built with the dimensions of 40 cm (length) × 120 cm (width) × 40 cm (height) (Supplementary Figure 2 and Figure 3A). The apparatus was made from Pexiglas and consisted of black walls and a white floor to provide contrast against the dark-colored Nile rats or B6 mice. Each of the three chambers (chamber A, the central chamber, and chamber B) measured 40 cm (length) × 40 cm (width) was separated by a black wall with a rectangular opening (8 cm width × 8cm height). We used two acrylic sheets as doors to manually control the access through these openings. Two cameras (DCS-8350LH, D-Link, Taipei, Taiwan) were positioned 50 cm above the center of each of the two adjacent chambers to capture animal movements comprehensively.

### Design of behavioral experiments

To evaluate the preference for WER, we adopted a modified protocol of the three-chambered sociability test^70^ by using still images as stimuli. Our study used still images instead of real animals as stimuli to eliminate the influence of olfactory cues that may affect the behavior of the test subjects. Before each test, the tested animals were acclimated in the experiment room for at least one hour. Each trial, consisting of a 10-min habituation phase followed by a 10-min testing phase, was investigated on one subject and was video-recorded. During the habituation phase, the animal was introduced to the center chamber and was allowed to freely explore the entire apparatus. After the habituation period, it was gently confined to the center chamber while doors to the side chambers were closed. We then placed still images corresponding to each experiment (as described below) on the side walls of Chamber A and Chamber B. The testing phase started once the doors were opened. After each trial, the floor and walls of the apparatus that the animal had contacted were cleaned with 70% ethanol. To reduce the number of animals used, some Nile rats or mice were used twice with at least a one-week interval between the two experiments to minimize memory effects.

Several sets of experiments on Nile rats were conducted as follows. The first experiment focused on the preference for WER on a male Nile rat’s face (Figure 3B). A still image of a male Nile rat with normal WER was placed on the side wall of Chamber A, while the same image, modified by Adobe Photoshop to mask both WER, was placed on the side wall of Chamber B. Thirty male and thirty female rats were evaluated in this setup. The second experiment tested the preference for WER on a female Nile rat’s face (Figure 3C), following the same procedure as the first experiment but instead using images of a female Nile rat. Similarly, thirty male and thirty female rats were evaluated. The third experiment examined the preference for WER on another species (Figure 3D), using images of an American red squirrel (*T. hudsonicus*), with one image showing normal WER and another with both WER masked. Fifteen male and fifteen female rats were tested. In the fourth experiment (top, Figure 3E), the preference for “asymmetrical facial pattern” was investigated by masking the left WER in a Nile rat image to create a complete but asymmetrical WER. This modified image was placed in Chamber A, while an image with both WER masked was placed in Chamber B. The fifth experiment explored the preference for a “symmetrical facial pattern” by masking the bottom half of the WER in a Nile rat image to create an incomplete but symmetrical WER (bottom, Figure 3E). The modified image was placed in chamber A, while an image with both WER masked was placed in chamber B. Fifteen male and fifteen female rats were evaluated in both the fourth and fifth experiments. To control for potential effects of stimulus novelty, we then repeated the fourth and fifth experiments, replacing the masked control image in Chamber B with an unmodified image showing normal WER. Fifteen male and fifteen female rats were tested in each repeated experiment. Results of these repeated tests are shown in Figure 3F.

A similar protocol was used to assess WER preference in B6 house mice. Test images depicted wild house mice with a natural agouti coat, either unaltered (image source: https://www.inaturalist.org/observations/300314476) or digitally modified by Adobe Photoshop to display WER. The image with WER was placed in Chamber A, and the unaltered image in Chamber B. Twenty male and twenty female mice were tested.

### Measuring social preference

Videos collected from the three-chambered sociability tests were analyzed by DeepLabCut version 2.3.8 to track animal pose and measured the time spent (in second) in each chamber. Side bias towards either chamber might occur during habituation due to confounding variables such as micro-environmental conditions, genetic background, or animal’s temperament and personality^71^. To determine the existence of side bias for each sex in each treatment, we compared the mean time spent in chamber A and chamber B during the habituation phase using the Wilcoxon signed-rank test.

The strength of preference (*SOP*) value for the stimulus in Chamber A relative to that in Chamber B was calculated for each trial using the formula:

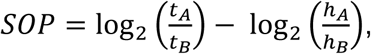

where *t_A_*, *t_B_* are the time spent in chamber A and B in testing phase, respectively, and *h_A_*, *h_B_* are the time spent in chamber A and B in habituation phase, respectively. Basically, *SOP* is the log ratio of time spent in chamber A to chamber B during the testing phase, adjusted for the animal’s baseline preference (the aforementioned side bias). A significantly *SOP* value greater than 0 (determined by a two-sided Wilcoxon signed-rank test) indicates that the animals showed a stronger interest in chamber A compared to chamber B. Conversely, an *SOP* value less than 0 suggests a preference for chamber B over chamber A. If *SOP* does not significantly differ from 0, the animals display no discernable preference for either chamber. In each treatment, differences of *SOP* were also compared between male and female animals using two-sided Mann-Whitney *U* test.

### Statistics and reproducibility

The statistical tools and parameters used to test phylogenetic signal, determine the best-fitting evolutionary models, reconstruct ancestral phenotypic states, and assess correlated evolution between focal traits are described in the corresponding sections above. Sample sizes for the behavioral tests used to generate violin plots in Figures 3 (Nile rats) and 4 (B6 house mice) are specified in the respective figure legends. Statistical analyses, including one-sample Wilcoxon signed-rank tests and two-sided Mann–Whitney *U* tests, were performed using the *wilcox.test* function in R (version 4.4.1; https://cran.r-project.org/). A *P*-value < 0.05 was considered statistically significant.

## Data availability

The phylogenetic tree in Newick format and phenotypic data of 601 rodent species can be found in Supplementary Data 1 and 2, respectively. The time spent in each chamber used to calculate *SOP* values can be found in Supplementary Data 3.

## Acknowledgments

We thank KC Hayes for providing Nile rat breeding pairs and advice in animal care. We thank Hon-Tsen Yu, Peng Shi, and Deepa Agashe for valuable discussions. This work was supported by an intramural grant from National Health Research Institutes, Taiwan, and the research grant (grant number 112-2311-B-400-003-MY3) from the National Science and Technology Council, Taiwan.

## Author contributions

BYL conceived the study; NHKL, SHL, CCC, and BYL designed trait evolution analyses; NHKL, MPW, SKC, and BYL designed behavioral experiments; NHKL and MPW conducted experiments; NHKL and MPW compiled and analyzed data; NHKL and BYL wrote the manuscript; SHL, CCC and SKC contributed to manuscript writing.

