## Supplementary Information for "White eye-rings coevolved with diurnal behaviors as a trait enhancing visual appeal in rodents"

**The Supplementary Information includes:**

- Supplementary Figures 1 to 2
- Supplementary Tables 1 to 7
- Supplementary Data 1 to 4 (separate files)
- Supplementary References

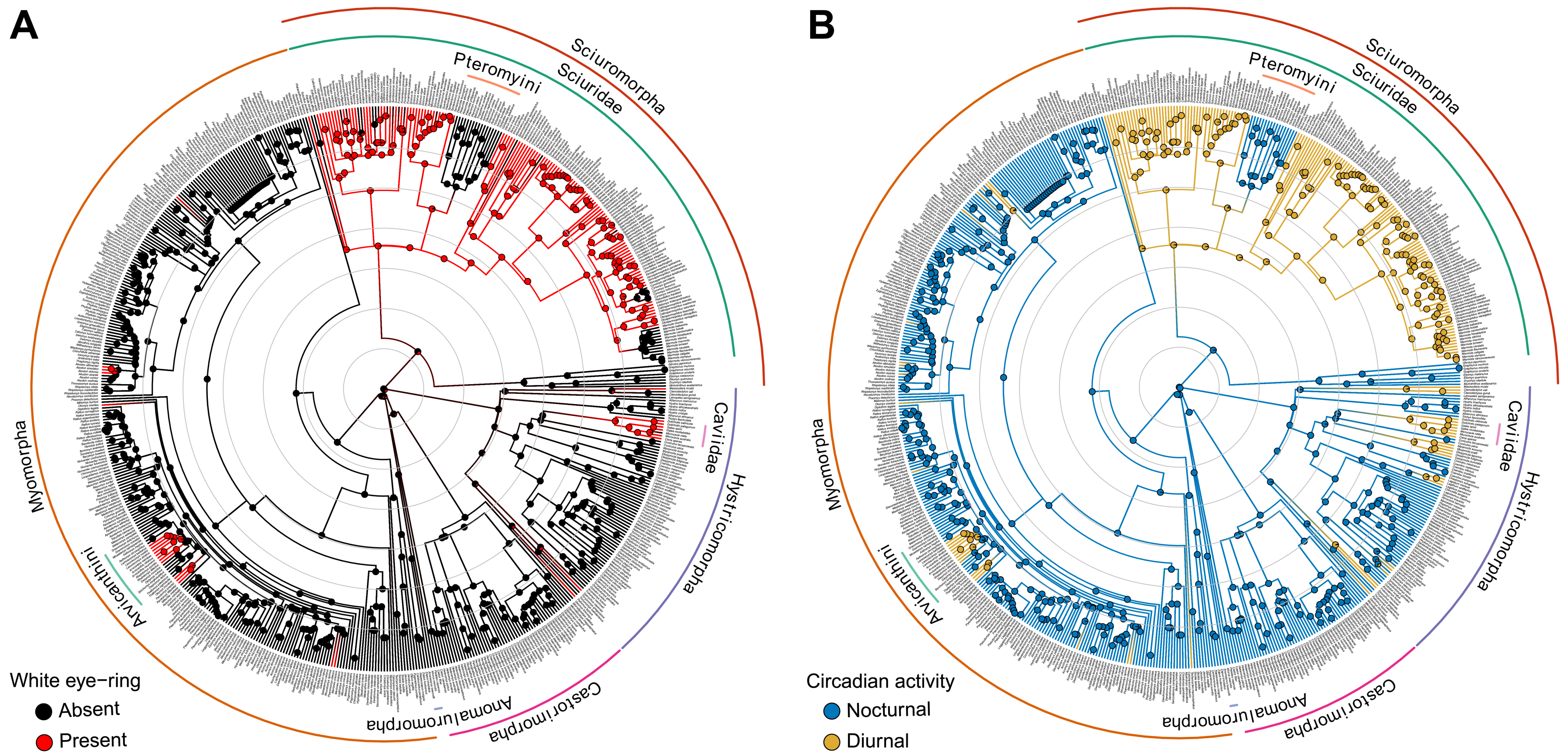


**Supplementary Figure 1.** Regenerated Figure 2A-B by adding species name information to each terminal tip and reconstructed posterior probabilities of the states to each node.

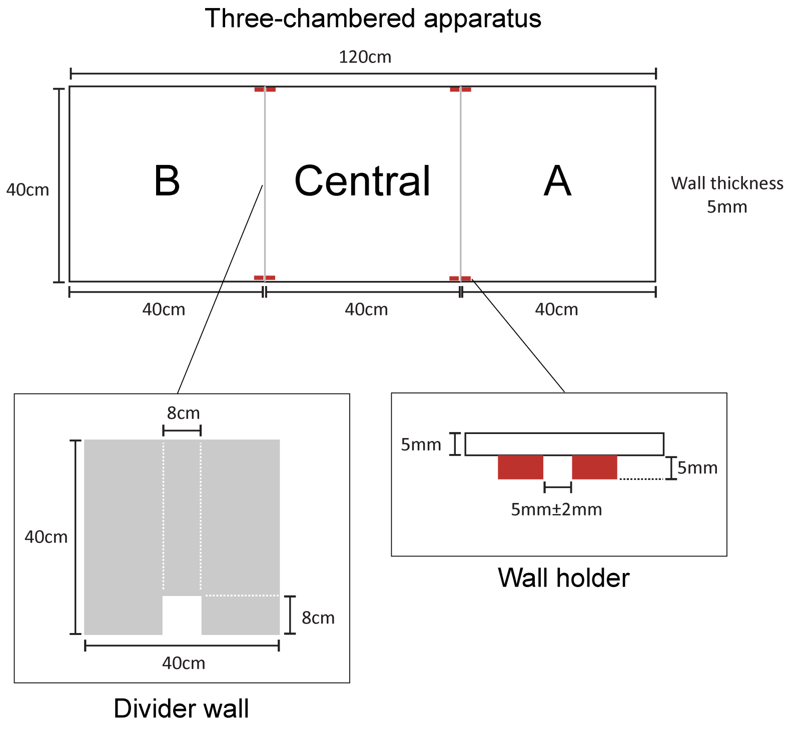


**Supplementary Figure 2.** The design of three-chambered apparatus used in the sociability test. The apparatus consists of three chambers of equal size (40 cm x 40 cm x 40 cm) including chamber A, central chamber, and chamber B. The chambers are divided by walls with an open doorway (8 cm x 8 cm).

**Supplementary Table 1.** Descriptions of phenotypic variables analyzed in this study.

| Traits | Definition | Values |
|---|---|---|
| White eye-ring (WER) | Light-colored pelage encircling the eyes with a distinct contrast and a well-defined boundary against the adjacent facial coloration of the animal | Absence / Presence |
| Non-WER markings | Additional white markings such as facial mask or stripes, body spots, blotches or stripes, black-and-white quills or tail | Absence / Presence |
| Circadian activity | Primary activity at night (nocturnal) or during daylight (diurnal) | Nocturnal / Diurnal |
| Habitat preference | | |
| Forest | Primary, secondary, deciduous, evergreen, montane, tropical, subtropical forest; woodlands; woody areas | Yes / No |
| Grassland | Meadows, brushland, shrubland, savanna, wetland, marshy area, steppe grasses | Yes / No |
| Desert | Desert, semi-desert, arid, coastal, sandy areas, rocky areas. | Yes / No |
| Social organization | Solitary or Pair living: Any species that exclusively spends time alone, with a partner, or with its offspring, and does not interact with other members of its species in groups of three or more<br>Social, group-living: Any species that occasionally gathers in groups of three or more individuals | Solitary / Group-living |

**Supplementary Table 2.** Count of species in the given taxon with the specified phenotypic trait.

| Taxa (total number of species) | WER | | Non-WER markings | | Circadian activity patterns | | Habitat preference | | | Social organization | |
|---|---|---|---|---|---|---|---|---|---|---|---|
| | Absent | Present | Absent | Present | Nocturnal | Diurnal | Forest (Yes) | Grassland (Yes) | Desert (Yes) | Solitary | Group-living |
| Rodentia (601) | 456 | 145 | 456 | 145 | 397 | 204 | 330 | 310 | 101 | 322 | 97 |
| Anomaluromorpha (3) | 3 | 0 | 2 | 1 | 3 | 0 | 2 | 1 | 1 | 1 | 2 |
| Castorimorpha (53) | 53 | 0 | 32 | 21 | 53 | 0 | 10 | 37 | 34 | 44 | 2 |
| Hystricomorpha (80) | 71 | 9 | 63 | 17 | 56 | 24 | 53 | 29 | 12 | 48 | 22 |
| Caviidae (7) | 1 | 6 | 0 | 7 | 0 | 7 | 2 | 7 | 0 | 4 | 3 |
| Myomorpha (289) | 269 | 20 | 264 | 25 | 260 | 29 | 152 | 164 | 40 | 144 | 41 |
| Muridae (139) | 123 | 16 | 118 | 21 | 121 | 18 | 66 | 84 | 22 | 51 | 34 |
| Arvicanthini (19) | 11 | 8 | 15 | 4 | 10 | 9 | 4 | 17 | 0 | 4 | 5 |
| Sciuromorpha (176) | 60 | 116 | 95 | 81 | 25 | 151 | 113 | 79 | 14 | 85 | 30 |
| Sciuridae (168) | 52 | 116 | 93 | 75 | 17 | 151 | 107 | 75 | 14 | 77 | 30 |
| Pteromyini (17) | 17 | 0 | 11 | 6 | 17 | 0 | 17 | 0 | 0 | 6 | 0 |

**Supplementary Table 3.** The divergence time (million years ago, MYA) and posterior probability (PP) of reconstructed ancestral states of WER and circadian activity pattern in different rodent lineages.

| Taxa | Divergence time (MYA) | PP of WER | | PP of circadian activity pattern | |
|---|---|---|---|---|---|
| | | Absence | Presence | Nocturnal | Diurnal |
| Rodentia | 70.222 | 0.780 | 0.220 | 1.000 | 0.000 |
| Anomaluromorpha | 48.538 | 0.916 | 0.084 | 1.000 | 0.000 |
| Castorimorpha | 63.474 | 0.877 | 0.123 | 1.000 | 0.000 |
| Hystricomorpha | 41.140 | 0.880 | 0.120 | 1.000 | 0.000 |
| Caviidae family | 11.920 | 0.009 | 0.991 | 0.013 | 0.987 |
| Myomorpha | 52.619 | 0.969 | 0.031 | 1.000 | 0.000 |
| Arvicanthini tribe | 8.535 | 0.990 | 0.010 | 1.000 | 0.000 |
| Diurnal branch of Arvicanthini | 7.349 | 0.026 | 0.974 | 0.036 | 0.964 |
| Sciuromorpha | 57.625 | 0.676 | 0.324 | 0.990 | 0.010 |
| Sciuridae family | 34.633 | 0.002 | 0.998 | 0.024 | 0.976 |
| Pteromyini tribe | 16.805 | 0.995 | 0.005 | 0.999 | 0.001 |

**Supplementary Table 4.** Testing for phylogenetic signals of white pelage and socioecological variables in rodents using *D*-statistic method.

| Traits | Estimated *D*-statistics | Brownian motion model (*P*-value, *D* = 0) | Phylogenetic randomness model (*P*-value, *D* = 1) |
|---|---|---|---|
| White eye-ring (WER) | -0.243 | 0.887 | 0.000 |
| Non-WER markings | 0.139 | 0.240 | 0.000 |
| Circadian activity | -0.316 | 0.953 | 0.000 |
| Habitat preference | | | |
| Forest | 0.165 | 0.183 | 0.000 |
| Grassland | 0.430 | 0.001 | 0.000 |
| Desert | 0.319 | 0.062 | 0.000 |
| Social organization | 0.495 | 0.003 | 0.000 |

**Supplementary Table 5.** Causal model testing for correlated evolution of WER and circadian activity patterns using Pagel's 1994 method.

| Dependent variables | Independent model | | Dependent model | | ΔAIC | *P*-value |
|---|---|---|---|---|---|---|
| | Log likelihood | AIC | Log likelihood | AIC | | |
| WER | -270.540 | 549.081 | -243.499 | 498.998 | 50.083 | 1.8E-12 |
| Circadian activity | -270.540 | 549.081 | -243.730 | 499.459 | 49.621 | 2.3E-12 |
| Interdependence | -270.540 | 549.081 | -237.373 | 490.746 | 58.335 | 1.4E-13 |

| Models | Akaike weights (AICw) |
|---|---|
| Independent | 2.1E-13 |
| WER depends on circadian activity | 0.0157 |
| Circadian activity depends on WER | 0.0125 |
| Interdependent WER and circadian activity | 0.9719 |

**Supplementary Table 6.** Phylogenetic logistic regression between the presence of WER and circadian activity patterns in Rodentia.

| Models | Slope | | | Intercept | | |
|---|---|---|---|---|---|---|
| | Estimates | SE | *P*-value | Estimates | SE | *P*-value |
| WER depends on Diurnalism [a] | 4.534 | 0.962 | 2.4E-06 | -5.377 | 1.078 | 6.0E-07 |
| Diurnalism depends on WER [b] | 1.991 | 0.463 | 1.7E-05 | 1.127 | 1.352 | 0.404 |

[a] Slope: diurnal; Intercept: nocturnal as reference
[b] Slope: presence of WER; Intercept: no WER as reference

**Supplementary Table 7.** Phylogenetic logistic regression between the presence of WER and confounding factors including the presence of non-WER markings, habitat preference, social organization.‡

| Models | | Coefficients | Estimated Log-odd ratio | SE | *P*-value |
|---|---|---|---|---|---|
| Non-WER markings | | Intercept (Absence as reference) | -1.758 | 0.856 | 0.040 |
| | | Slope (Presence) | 0.302 | 0.263 | 0.249 |
| Habitat preference | | | | | |
| | Forest | Intercept (No as reference) | -1.629 | 0.763 | 0.033 |
| | | Slope (Yes) | 0.053 | 0.153 | 0.728 |
| | Grassland | Intercept (No as reference) | -1.641 | 0.961 | 0.088 |
| | | Slope (Yes) | 0.201 | 0.154 | 0.192 |
| | Desert | Intercept (No as reference) | -1.535 | 0.785 | 0.050 |
| | | Slope (Yes) | -0.160 | 0.225 | 0.477 |
| Social organization | | Intercept (Solitary as reference) | -1.243 | 1.122 | 0.268 |
| | | Slope (Group-living) | -0.365 | 0.208 | 0.079 |

‡All phylogenetic logistic regression analyses in this study used the presence of WER markings as the dependent variable. The slope coefficient in each logistic regression represents the log-odds ratio of the presence of WER. For non-WER markings, habitat preference, and social organization, the presence of the respective trait (markings, specific habitat, or group-living) was assigned as the slope, while the other value of the variable served as the reference. Note that while the choice of reference category in binary logistic regression does not affect the model's overall fit or statistical significance, it does influence the interpretation of the coefficients. In this study, the specific interpretations of the slopes are based on the chosen reference categories as outlined above.

**Supplementary Data 1 (separate file).** The phylogenetic tree of the 601 rodent species examined in this study, provided in Newick format.

**Supplementary Data 2 (separate file).** Taxonomy and phenotypes in 601 rodent species. The websites linked to image sources used to determine the presence or absence of WER, as well as references for habitat preferences and social structure, are provided.

**Supplementary Data 3 (separate file).** Time spent (in second) and strength of preference (*SOP*) values of Nile rat subjects during the three-chambered behavioral tests.

**Supplementary Data 4 (separate file).** Time spent (in second) and strength of preference (*SOP*) values of house mouse subjects during the three-chambered behavioral tests.
